# The MarR Family Transcription Factor SlyA Senses Iron and Respiratory Status in Enteric Bacteria

**DOI:** 10.1101/2025.04.07.647610

**Authors:** W. Ryan Will, Ferric C. Fang

## Abstract

SlyA and its homologs are conserved essential transcription factors in enteric bacteria, including *Salmonella enterica* and *Yersinia pestis*, in which they upregulate horizontally-acquired virulence genes. As members of the MarR family of transcription factors, they possess a small molecule binding pocket that allows the protein to undergo a conformational change upon ligand interaction that abrogates DNA binding. Although the original discovery of MarR was based on its ability to recognize xenobiotic compounds and promote their efflux, the conservation of the MarR family throughout Bacteria and Archaea suggests a more general function. Because SlyA is known to bind xenobiotic aromatic carboxylates, we performed a targeted analysis of aromatic metabolic genes in *S*. Typhimurium to identify potential endogenous ligands, including genes involved in essential cellular processes including iron metabolism, respiration, and aromatic amino acid and folate biosynthesis. We found that SlyA is promiscuously inhibited by multiple aromatic carboxylates including 2,3-dihydroxybenzoate, a precursor of iron-scavenging catecholate siderophores, and 4-hydroxybenzoate, a precursor of quinone-based electron-carriers, which allows it to sense changes in iron availability, respiration and growth on succinate. We suggest that SlyA and other MarR proteins sense bacterial metabolic status via the flux of aromatic carboxylates in biosynthetic pathways, allowing SlyA to function as a counter-silencer of horizontally-acquired genes that is exquisitely responsive to the metabolic state of the cell.

**SIGNIFICANCE:** MarR proteins comprise an ancient group of transcription factors that emerged before the divergence of Archaea and Bacteria. First identified as regulators of antibiotic efflux, they have also been suggested to sense and regulate concentrations of endogenous intracellular metabolites, but such metabolites have not previously been identified. Here we show that SlyA, a conserved and essential MarR family virulence gene regulator in enteric pathogens, binds to and is inhibited by aromatic metabolites required for many essential cellular processes. Our findings show how SlyA integrates metabolic status into bacterial transcriptional networks that control both molecular efflux and virulence.

## INTRODUCTION

All living organisms respond to internal physiological and external environmental cues to optimize growth and mitigate stress through systems that sense and integrate these signals to modulate gene expression. The MarR family of transcription factors (MFTFs) is an ancient protein lineage that has been hypothesized to promote cellular homeostasis by detecting toxic metabolites and regulating the expression of small molecule efflux pumps (1). MFTFs comprise one of the earliest helix-turn-helix (HTH) containing protein families in nature, pre-dating the divergence of Archaea and Bacteria (2, 3). MFTFs are ubiquitous in bacteria, with the average genome encoding ∼7 unique family members (4). Members of this family typically include: (1) a winged helix-turn-helix (wHTH) domain, (2) a compact globular structure, (3) small molecule binding sites that promote allosteric inhibition, and (4) frequent genetic linkage to efflux pumps (5). Although MFTFs originally evolved as repressors, many have also acquired the ability to upregulate gene expression by acting as counter-silencers (5). Although the reasons for their ubiquity and conservation throughout multiple domains of life is unclear, the ligand binding domains and linkage to small molecule efflux support the notion they may have first emerged as primordial metabolic sensors (1).

MarR was the first MFTF to be identified, due to its role as a repressor of multiple antibiotic resistance in *Escherichia coli* (6). Although it is often considered to be a prototype for the MFTF family, MarR is limited to a few enteric bacterial species, in which it is expressed at low levels and is only functionally, but not physically, linked to efflux pump genes. Therefore, MarR may not be representative of MFTFs in general. Another MFTF called SlyA is broadly conserved and highly expressed in *Enterobacterales* (5). SlyA exhibits the classic characteristics of MFTFs, including allosteric inhibition by small molecules (7) and direct linkage to an efflux pump (5), suggesting that it may be more typical of MFTF family members.

SlyA is also of interest for its established role as a regulator of genes required for bacterial virulence and for its important role in facilitating bacterial evolution (8). Bacteria evolve predominantly by horizontal gene transfer, which allows species to acquire new traits, such as virulence and antibiotic resistance, through a single transfer event (9). Newly acquired genes must be successfully integrated into existing regulatory networks to achieve physiologically appropriate gene expression, which is achieved by the balance between the mechanisms of xenogeneic silencing and counter-silencing (8). Xenogeneic silencers recognize horizontally-acquired DNA with increased AT content, oligomerizing along adjacent regions of DNA to silence gene expression (10). The histone-like nucleoid associated (H-NS) protein was the first nucleoid-associated protein shown to function as a xenogeneic silencer in the *Enterobacterales* (11). Xenogeneic silencing appears to be a widespread paradigm in bacteria, and genetically distinct but functionally analogous proteins have been subsequently identified in other species, including MucR in α-Proteobacteria (12, 13), Lsr2 in *Actinomycetes* (14, 15), MvaT in *Pseudomonas spp*. (16, 17) and Rok in *Bacillus spp*. (18). DNA-binding proteins with greater sequence specificity and binding affinity, exemplified by SlyA in enteric bacteria (5), compete with silencing proteins to disrupt silencing complexes and allow transcription. The involvement of specific counter-silencers allows horizontally-acquired genes to be incorporated into existing regulatory networks and expressed under appropriate environmental conditions. Recent studies suggest that even in species lacking H-NS homologs, SlyA-related MFTFs are able to function as counter-silencers (13), suggesting that these proteins possess traits that predispose them to this function, which plays a fundamental role in the evolution of bacterial regulatory networks. One possible explanation for their preferred role as counter-silencers is the ability to sense the metabolic status of the cell by interacting with specific metabolites.

However, the identity of putative ubiquitous endogenous ligands to account for the conservation of MFTFs as counter-silencers has not been established. The aromatic carboxylate, salicylate, was shown to inhibit MarR when a panel of compounds known to induce multiple antibiotic resistance was screened for *marR*-dependent activity (6). However, salicylate is not a conserved metabolite across bacterial species. Other aromatic carboxylates such as hydroxycinnamic acid have also been identified as potential ligands (19) but are similarly not conserved. A subfamily of MFTFs, including OhrR of *Bacillus subtilis* (20, 21) and possibly MarR itself (22), are subject to regulation by cysteine oxidation, which induces conformational changes and subsequent dissociation from DNA, but genetic analyses have shown that this mechanism is not conserved in all MFTFs, including SlyA (23).

The objective of the present study was to identify endogenous ligands of SlyA in the model enteric organism *Salmonella enterica* serovar Typhimurium, which might account for the ubiquity of SlyA and other MFTFs throughout the prokaryotes. *S*. Typhimurium is an important intracellular pathogen in humans and many animal species, causing both acute and chronic disease (24). *S*. Typhimurium virulence is primarily dependent on two genomic islands, Salmonella Pathogenicity Islands 1 and 2 (SPI-1 and -2), each encoding a type III secretion system and multiple effectors that act on host cell targets (25-28). SPI-1 is required for the invasion of non-phagocytic host cells, whereas SPI-2 is required for survival in macrophages within a modified phagosome termed the Salmonella-containing vacuole (SCV) (29). SlyA counter-silences SPI-2 genes in *S*. Typhimurium (7, 23), facilitating both replication in the SCV and systemic infection, but SlyA homologs also counter-silence a variety of virulence genes in other bacterial pathogens, including *Yersinia spp*. (30), *Dickeya spp*. (31), *Shigella flexneri* (32), and *Klebsiella pneumoniae* (33). The conservation of SlyA in the *Enterobacterales* suggests that its endogenous physiological ligands are likely to be central metabolites relevant to both virulence and antibiotic resistance, which could sensitize SlyA, and other MFTFs to the metabolic status of the cell. In the present study, we conducted a targeted screen of aromatic carboxylate metabolic genes in *S*. Typhimurium and found that SlyA responds to the metabolic flux of aromatic carboxylates. We next characterized 4-hydroxybenzoate (4-HB) and 2,3-dihydroxybenzoate (2,3-DHB) as endogenous ligands for SlyA, which sensitize virulence and resistance genes to the respiratory and iron nutritional status of the cell, identifying SlyA as a novel iron-responsive regulator that is independent of both the conserved Ferric uptake regulator (Fur) and its associated small regulatory RNA, RyhB (34). Collectively, our observations have identified endogenous physiological ligands for SlyA, supporting the hypothesis that MFTFs have evolved to sense metabolic status via aromatic carboxylates, which facilitates their important role as counter-silencers of gene expression.

## RESULTS

### Inhibition of efflux inhibits SlyA counter-silencing

We hypothesized that if SlyA or other MFTFs respond to endogenous ligands, then disruption of trans-membrane efflux might result in ligand accumulation and SlyA inhibition. Bacteria typically encode multiple efflux pumps, but the Resistance-Nodulation-Division (RND) family of pumps is the most significant contributor to antimicrobial resistance, and RND pumps in Gram-negative bacteria share a common subunit called TolC (35). A null mutation of *tolC* was constructed, and RNA-Seq analyses of wildtype, *slyA*, and *tolC* mutant *S*. Typhimurium strains in minimal medium with low Mg^2+^ concentrations was performed. These conditions mimic the SCV, resulting in expression of the PhoPQ two-component signal transduction system and SlyA (36), which functions cooperatively with PhoPQ. SlyA was found to regulate more than 133 genes (Fig. 1a,c; Dataset S1), many of which were reported previously (7). The majority of the SlyA regulon is up-regulated, and the genes most strongly up-regulated by SlyA were predominantly from SPI-2, including the model counter-silenced gene, *pagC*. However, core genes were also repressed by SlyA, including several involved in methionine biosynthesis, the *ydhIJK* efflux pump operon, which is transcribed divergently from *slyA*, and the *marRAB* operon. Mutation of *tolC* altered the expression of more than 553 genes (Fig. 1b,c; Dataset S2), comprising approximately 11% of the *S*. Typhimurium genome. Greater than half of the SlyA regulon (69/133, 52%), including 54.5% of up-regulated genes (43/79), were co-regulated by TolC. This fraction is even greater when considering the most strongly up-regulated genes (14/20), which are most likely to be directly counter-silenced, supporting the hypothesis that TolC exports an endogenous inhibitory ligand of SlyA. TolC was also observed to regulate many virulence genes outside of SPI-2, including *sopA, B, D, D2* and *E2, hilA* and *D*, and *invFGEABC* and *invH*, suggesting that additional virulence regulators are affected by efflux. Transport genes in the core genome were broadly influenced, including porins (*ompC, F*, and *N*), a dipeptide permease (*dppABCDF*), multidrug transporters (*mdtABC*; *acrD*), and additional specific and putative transporters. Ribosomal genes were modestly but broadly down-regulated. Genes exhibiting altered expression included those associated with tryptophan and enterobactin biosynthesis, processes with aromatic intermediates. The *trpS2, mtr*, and *trpABCDE* genes were significantly down-regulated in a *tolC* mutant, while *entCEBAH* genes associated with enterobactin biosynthesis were significantly up-regulated. Both enterobactin and tryptophan share the common aromatic precursor chorismate, which is involved in the synthesis of multiple aromatic compounds.

**Figure 1.**
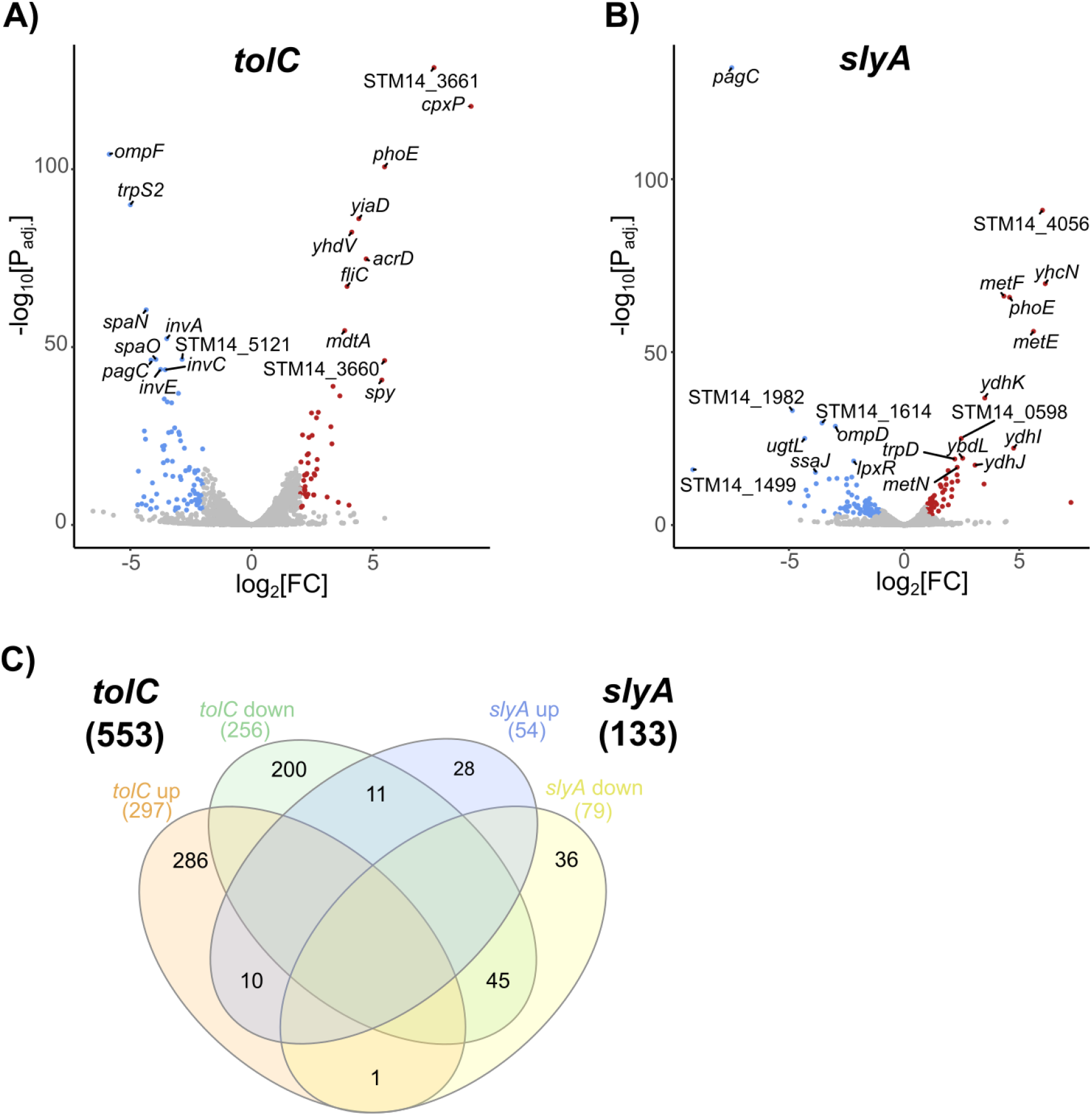
Transcriptomic analysis of *tolC* and *slyA*. Wildtype 14028s or isogenic *tolC*, or *slyA* mutant strains were grown to late exponential phase in LB medium before inducing virulence gene expression for 1 hr in N-minimal medium containing 10 µM MgSO_4_. Volcano plots of the *tolC* (A) and *slyA* (B) regulons are shown. Genes significantly up-regulated (≥2 fold-change; P_adj._≤ 0.05) in a mutant strain are indicated by red dots, whereas genes that are significantly down-regulated are indicated by blue dots. Log_2_ fold-change is indicated on the x-axis, whereas -log_10_ P_adj_ is indicated on the y-axis. A Venn diagram of the two regulons (C) shows that the majority of SlyA regulated genes (∼52%) are co-regulated by TolC.

### Disruption of ubiquinone or enterobactin synthesis inhibits SlyA counter-silencing

Many MFTFs bind aromatic carboxylates such as salicylate (6, 19). We hypothesized that an endogenous ligand was also likely to be an aromatic carboxylate, compounds that are synthesized from phosphorenolpyruvate and erythrose-4-phosphate in the pentose phosphate pathway, which produces the aromatic carboxylate, dehydroquinic acid (37) (Fig. 2a). Dehydroquinic acid undergoes seven distinct enzymatic reactions to become shikimate and eventually chorismate, which is the final universal aromatic intermediate (38). Chorismate is a metabolic hub and precursor for several pathways, including the synthesis of the aromatic amino acids, tetrahydrofolate, quinones, and iron-scavenging siderophores. The isoprenoid quinones, including both the primary Gram-negative membrane electron carrier ubiquinone and the alternative napthoquinones, such as menaquinone (39), are derived from chorismate and function as electron carriers in all bacteria, playing a critical role in respiration and the generation of proton motive force. Catecholate siderophores such as enterobactin are responsible for scavenging iron in limiting environments (40) and are tightly coupled to electron transfer and respiration, as many of the enzymes in these processes require iron as a co-factor in iron-sulfur clusters (41, 42). Given the central role of chorismate and its related metabolites in bacterial metabolism, we systematically mutated each branch of aromatic carboxylate metabolism upstream and downstream of chorismate and examined the effects on the expression of a *pagC* reporter under up-regulating, low Mg^2+^ conditions. Under these conditions, the mutation of any enzyme upstream of a ligand in the pathway was expected to result in the depletion of that ligand, resulting in an increase in SlyA-mediated counter-silencing and *pagC* expression. Conversely, the mutation of enzymes acting downstream of a putative ligand was anticipated to result in ligand accumulation, thereby decreasing counter-silencing and *pagC* expression. In addition, it is possible that blocking a single chorismate-dependent pathway might result in enhanced flux down another branch of chorismate metabolism.

**Figure 2.**
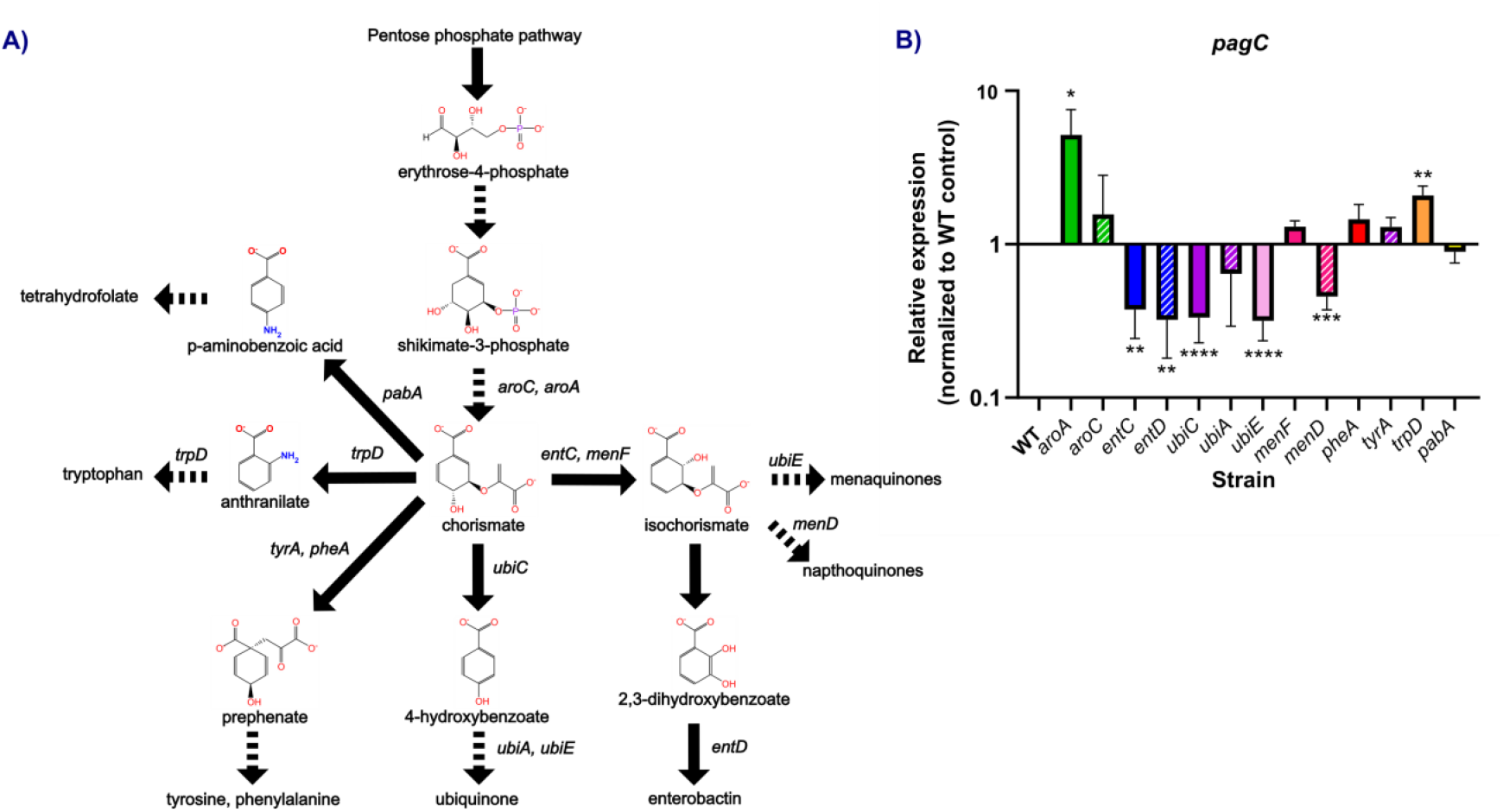
Aromatic carboxylate metabolism regulates virulence gene expression. Aromatic carboxylate compounds in *S*. Typhimurium are synthesized from erythrose-4-phosphate, which is produced from the pentose phosphate pathway (A). Erythrose-4-phosphate is converted to chorismate, the final universal intermediate in aromatic carboxylate metabolism, which is a precursor for the biosynthesis of folate, aromatic amino acids, quinones, and enterobactin. Biosynthetic genes of interest are indicated next to the reactions catalyzed by the corresponding enzyme. Solid arrows indicate a single enzymatic step, while dashed arrows indicate multiple enzymatic steps. To determine whether aromatic carboxylate metabolism influences the expression of the SlyA-dependent *pagC* reporter gene, each of the indicated enzymes were mutated and *pagC* expression determined after growth in N-minimal medium containing 10 µM MgSO_4_, which mimics conditions associated with the SCV. Results are normalized to *rpoD* and indicate mean ± S.D. (n ≥ 3). Asterisks (*, **, ***, ****) indicate P-values ≤ 0.05, 0.01, 0.0005 and 0.0001, respectively.

Mutation of either *aroB*, which is required for the synthesis of dehydroquinate, or *aroC*, which synthesizes chorismate, resulted in an increase in *pagC* reporter expression, indicating that endogenous ligands are synthesized downstream of these enzymes (Fig. 2b). Chorismate can subsequently follow one of five metabolic paths, (a) conversion to ρ-aminobenzoic acid for folate synthesis, (b) conversion to anthranilate for tryptophan synthesis, (c) conversion to prephenate for tyrosine and phenylalanine synthesis, (d) conversion to 4-HB for ubiquinone synthesis, and (e) conversion to isochorismate for either enterobactin or napthoquinone synthesis. Mutation of *pabA* in the folate biosynthetic pathway had no detectable effect on *pagC* expression, nor did *tyrA* and *pheA*, required for tyrosine and phenylalanine biosynthesis, respectively. Mutation of *trpD*, which converts chorismate to anthranilate and then to 5-phosphoribosyl-anthranilate in consecutive reactions, resulted in a modest two-fold increase in *pagC* expression, suggesting that anthranilate, 5-phosphoribosyl-anthranilate, or another intermediate immediately downstream might serve as a ligand. With the exception of anthranilate, many of these compounds have extensive side groups, making them unlikely candidates for interaction with the compact ligand binding sites of SlyA (23). However, mutation of genes in the ubiquinone biosynthetic pathway, including *ubiC, ubiA*, and *ubiE*, each decreased *pagC* expression, indicating that SlyA is inhibited by one or more ubiquinone precursors. Chorismate is converted to 4-HB by UbiC (43) and then isoprenylated by UbiA (44), with the subsequent metabolites associated with the membrane and unavailable for interaction with cytoplasmic SlyA. Chorismate and 4-HB were further investigated as candidate ligands.

Mutation of the genes *entC* and *entD* in the enterobactin biosynthesis pathway also decreased *pagC* expression, indicating that this pathway is also sensed by SlyA. EntC is a chorismate mutase, responsible for converting chorismate to isochorismate. Isochorismate is then metabolized to 2,3-DHB, which the EntBDEF enterobactin synthase metabolizes to enterobactin (45, 46). We concluded that isochorismate and 2,3-DHB were also potential ligands. Although mutation of *menF*, encoding the chorismate mutase associated with the alternative napthoquinone biosynthetic pathways (47), did not affect *pagC* expression, EntC may be able to complement the loss of MenF. Mutation of *menD*, which converts isochorismate to 2-succinyl-5-enolpyruvyl-6-hydroxycyclohex-3-ene-1-carboxylate (48), also decreased *pagC* expression, suggesting that SlyA is in fact sensitive to perturbations of napthoquinone biosynthesis. Collectively, these observations suggest that SlyA is promiscuously inhibited by multiple endogenous aromatic ligands, including anthranilate, chorismate, isochorismate, 4-HB, and 2,3-DHB. These data also suggest that SlyA is sensitive to both iron availability and the closely linked respiratory status of the cell via changes in metabolic flux through these pathways.

To confirm that observed changes in gene expression were directly attributable to the accumulation of specific metabolites, metabolites were directly added to *S*. Typhimurium cultures carrying a *pagC-egfp* transcriptional reporter (Fig. S1.). As both iron metabolism and respiration are critical to the infectious process (42, 49), we focused on 4-HB and 2,3-DHB, and their precursors shikimate and dehydroquinic acid, each of which are commercially available in highly purified form. We also examined chorismate, which is commercially available but contaminated with related aromatic compounds such as 4-HB and 2,3-DHB. Cultures were grown to exponential phase and induced with low Mg^2+^ (10 µM) minimal medium before incubation in the presence of increasing concentrations of candidate ligands. Salicylate, included as a positive control, significantly inhibited reporter activity in a dose-dependent manner. However, neither of the aromatic chorismate precursors, dehydroquinic acid and shikimic acid, repressed reporter activity. Chorismate inhibited reporter activity only at the highest concentrations (>1 mM), suggesting that it has weak inhibitory activity at best. However, the addition of either 4-HB or 2,3-DHB resulted in a significant dose-dependent decrease in reporter activity starting at concentrations as low as 50 µM. This indicates that either intermediate can act as an inhibitory ligand to provide a link between respiration, iron nutritional status and SlyA-dependent gene expression.

### 4-hydroxybenzoate and 2,3-dihydroxybenzoate allosterically inhibit SlyA

To confirm that SlyA directly binds 4-HB and 2,3-DHB, isothermal titration calorimetry (ITC) was performed (Fig. 3). Chorismate was not included in these analyses due to its low purity. Aliquots of 4-HB (Fig. 3a) or 2,3-DHB (Fig. 3b) were added to purified SlyA and the resulting reaction heat measured to determine the thermodynamic properties of each binding reaction. Salicylate, used as a positive control, bound to SlyA with a K_*D*_ of 36 µM and a stoichiometry of ∼1 (Fig. S2), differing from our previously published structural analyses of the SlyA-salicylate complex, which identified two binding sites per subunit (23). However, mutational analysis in that study suggested only one site was required for allosteric inhibition by salicylate, suggesting that salicylate binding at the second site was a non-specific artifact. The lowest affinity of the three tested compounds was exhibited by 4-HB, which had a calculated K_*D*_ of ∼100 µM and predicted stoichiometry of ∼2, indicating that each SlyA subunit binds two 4-HB molecules. Notably, previous metabolic analyses in *E. coli* have determined intracellular concentrations of 4-HB to range from 52 µM to 787 µM, depending on growth conditions (50). The calculated K_*D*_ of 2,3-DHB was ∼57 µM with a predicted stoichiometry of only ∼1. In *E. coli*, Intracellular concentrations of 2,3-DHB have been determined to range from 138 µM to 414 µM, and both 4-HB and 2,3-DHB are actively exported into the extracellular medium (50). These measurements indicate that both compounds are present at intracellular concentrations that could influence SlyA activity, and disruption of efflux pump function, as in a *tolC* mutant strain, would be anticipated to affect the intracellular concentrations of both compounds.

**Figure 3.**
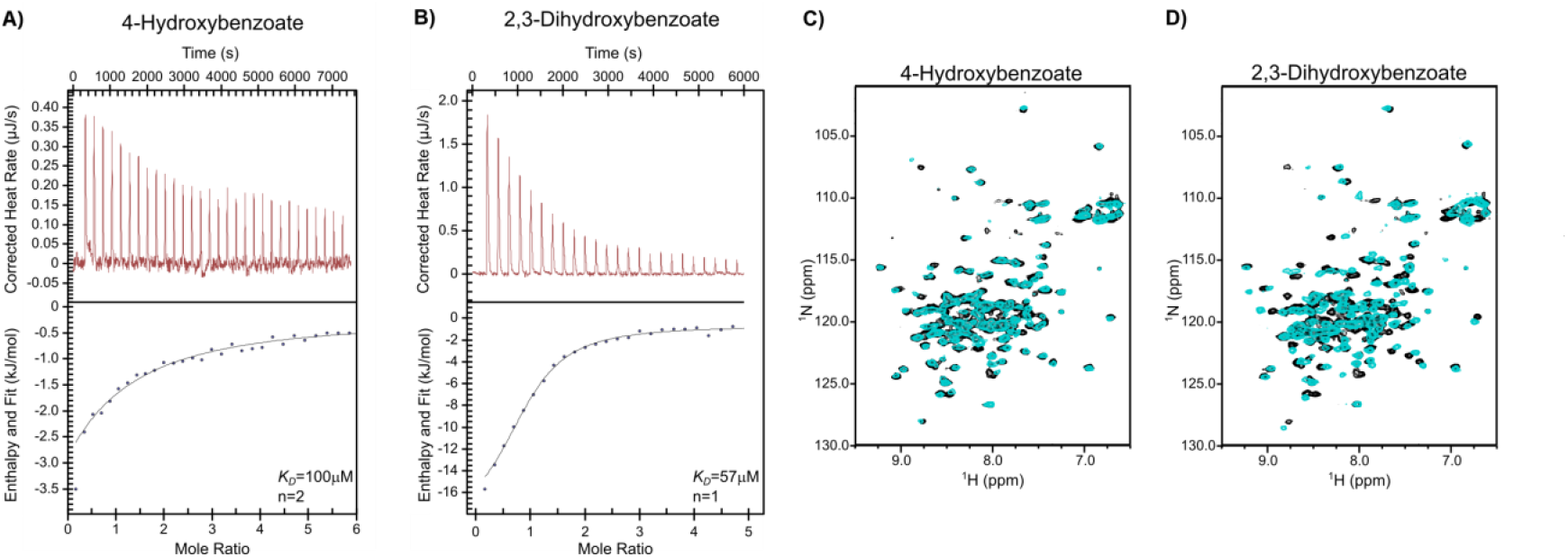
4-hydroxybenzoate and 2,3-dihydroxybenzoate directly bind SlyA and induce allosteric changes. Isothermal titration calorimetry was performed with SlyA and pure 4-HB (A) or 2,3-DHB (B). Binding reactions were performed in triplicate at 10ºC with aliquots of ligand injected at 4 min intervals. Upper panels are thermographs, measuring heat generated following each ligand injection. The enthalpy and stoichiometry of each injection are shown in the bottom plots. Each experiment was performed three times, and representative data are shown. To determine if ligand binding induces conformational changes, ^1^H,^15^N-HSQC NMR spectroscopy was performed on uniformly labelled ^15^N-SlyA in the presence (cyan) or absence (black) of 4-HB (C) or 2,3-DHB (D). Ligands were added to SlyA at a 4:1 molar ratio (1.2 mM:300 µM) and incubated at 35ºC. Spectra of SlyA-ligand complexes at different molar ratios are shown in Fig. S3.

We previously demonstrated that salicylate binding causes the recognition helix of SlyA to pivot out of register with the major helix of the DNA duplex to prevent DNA binding (23). We corroborated this structural change by nuclear magnetic resonance (NMR) spectrometry of apo- and salicylate-bound SlyA, observing a significant change in the NMR spectra upon salicylate binding (Fig. S2b). To determine whether 4-HB and 2,3-DHB induce structural changes similar to those observed with salicylate, we performed ^1^H,^15^N heteronuclear single quantum coherence (HSQC) NMR spectrometry on SlyA in the presence or absence of each compound (Fig. 3C,D). Binding by 2,3-DHB induced a shift similar to that observed for salicylate, indicating that the 2,3-DHB-SlyA complex was also allosterically inhibited. Although 4-HB induced a weaker shift than either salicylate or 2,3-DHB, a similar pattern was observed. The decreased magnitude of shifting may be attributable to the lower affinity of SlyA for 4-HB, and titration of SlyA with higher concentrations of 4-HB did somewhat increase the observed shift, albeit not to the degree observed with the other compounds (Fig. S3). Apo-SlyA exhibited a relatively high degree of variation or overlap among individual resonances, but each of the three ligands improved the resolution of individual resonances, with 2,3-DHB and salicylate having the most pronounced effect. This suggests that apo-SlyA may be disordered, existing in multiple conformational states, as previously suggested (23), but the addition of a ligand stabilizes the protein in a conformation that is no longer compatible with DNA binding. Together, these data indicate that both 4-HB and 2,3-DHB can directly bind and inhibit SlyA, and 2,3-DHB is the most effective endogenous allosteric inhibitor.

### SlyA counter-silencing is modulated by iron deprivation and growth on succinate

To establish the physiological relevance of aromatic carboxylate-mediated inhibition of SlyA, we examined SlyA-mediated counter-silencing of *pagC* under conditions likely to perturb aromatic metabolism. Enterobactin biosynthetic genes are repressed by the iron-dependent regulator Fur under iron-replete conditions (51, 52). However, when iron is limiting, Fur dissociates from DNA, allowing the transcription of enterobactin biosynthetic genes, which leads to the metabolism of chorismate. Intracellular concentrations of the aromatic precursors of enterobactin, including chorismate and 2,3-dihydroxybenzoate, increase as enterobactin biosynthesis is accelerated to allow the scavenging of sufficient iron for growth. To deplete bacterial cultures of iron and perturb this circuit, we added increasing concentrations of the iron chelator 2,2-dipyridyl under SlyA-inducing conditions, observing a dose-dependent effect on *pagC* expression (Fig. 4a). As the concentration of 2,2-dipyridyl increased, the expression of *pagC* decreased, most likely due to an increase in intracellular concentration of 2,3-DHB and other enterobactin precursors. Therefore, we established that SlyA-mediated counter-silencing is responsive to iron availability. To confirm that 2,2-dipyridyl-mediated inhibition was occurring via the induction of enterobactin synthesis, we examined the effects of iron chelation in *aroC* and *entC* mutant bacteria carrying a *pagC-egfp* reporter (Fig. 4b). The *aroC* mutant strain was sensitive to *2,2*-dipyridyl, although overall fluorescence and sensitivity decreased slightly. However, the 2,2-dipyridyl response was completely abrogated in the *entC* mutant strain, confirming that SlyA responds to iron availability via the enterobactin biosynthetic pathway.

**Figure 4.**
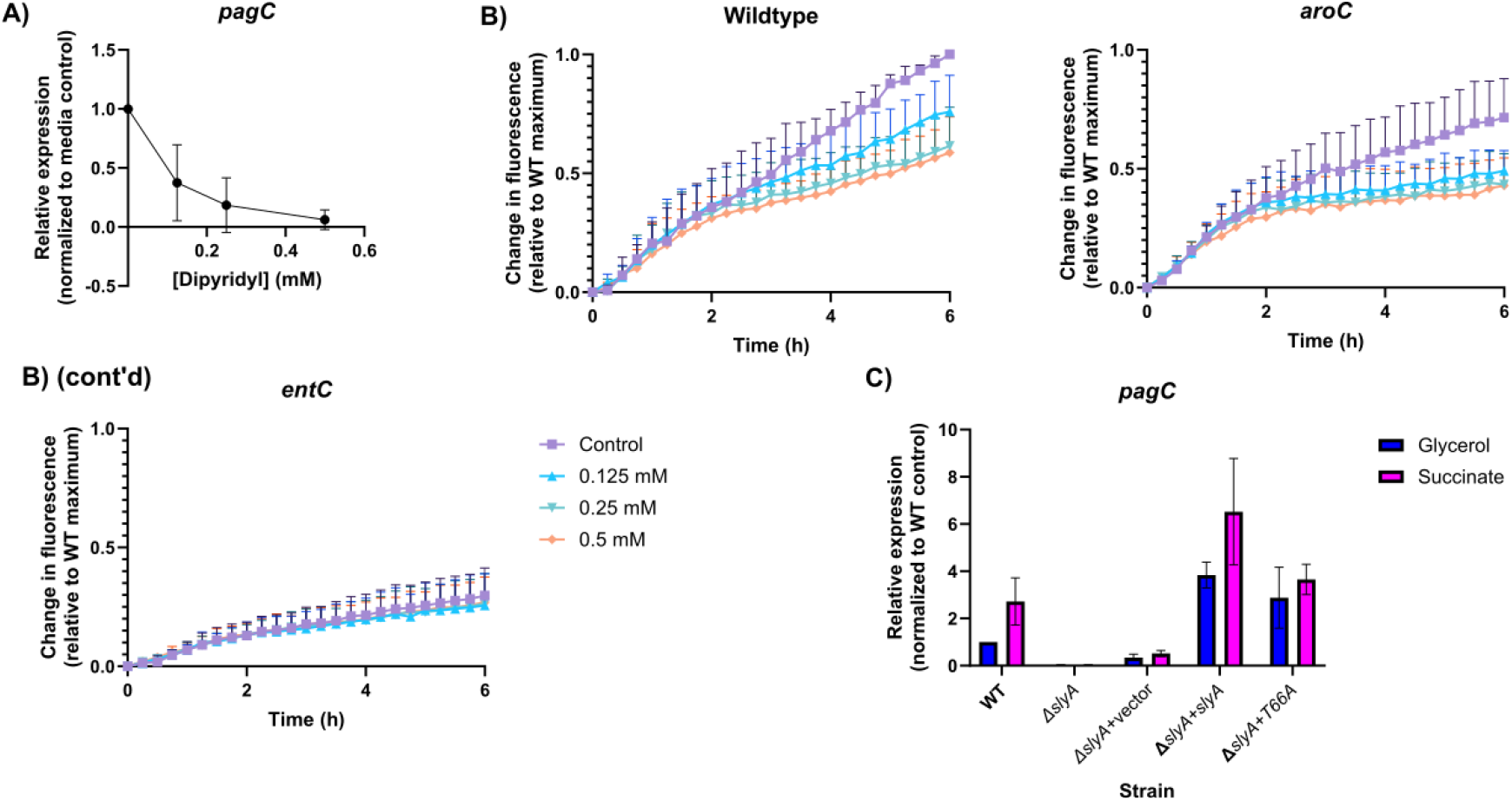
SlyA is regulated by iron availability and respiratory status. Expression of the SlyA-dependent reporter gene *pagC* was measured in cultures grown in N-minimal medium (10 µM MgSO_4_) in the presence of increasing concentrations of the iron chelator 2,2-dipyridyl. Data represent the mean ± S.D. (n=3). To determine whether SlyA-2,3-dihydroxybenzoate interactions were responsible for inhibition by 2,2-dipyridyl, wild-type, *aroC* mutant (blocking the production of chorismate) and *entC* mutant (blocking the production of 2,3-dihydroxybenzoate) strains containing the *pagC-egfp* reporter plasmid pRW79 were grown in N-minimal medium (10 µM MgSO_4_) with increasing concentrations of 2,2-dipyridyl. Results are normalized to the maximal fluorescence of the wild-type strain. Data represent the mean + S.D. (n=3). To measure the influence of respiration on SlyA activity, *S*. Typhimurium cultures were grown in LPM containing either glycerol or succinate as the sole carbon source. A *slyA* mutant strain was complemented *in trans* with either wild-type *slyA* or the T66A mutant allele, which we renders SlyA insensitive to ligand binding (23). Data represent the mean ± S.D. (n=3).

Other environmental changes not directly related to iron availability might also perturb these pathways, as the infectious process is associated with many metabolic changes in both the host cell and the bacterium. In the case of *S*. Typhimurium, uptake by macrophages induces a shift in host metabolism from oxidative phosphorylation to glycolysis and the accumulation of the tricarboxylic acid cycle intermediate succinate (53, 54). The increase in succinate has multiple implications for *Salmonella*, as it serves as both a carbon source and an electron donor, directly contributing electrons to ubiquinone via succinate dehydrogenase, to produce ubiquinol. A surge in succinate production has been shown to promote virulence gene expression, particularly those of SPI-2, which has been termed “succinate induction.” Although a mechanistic explanation for succinate induction has not been determined, transcriptomic analysis of cultures utilizing succinate as a sole carbon source has demonstrated decreased expression of several genes involved in chorismate metabolism, including critical enzymes in both the enterobactin (*entC* and *F*) and ubiquinol (*ubiC*) biosynthetic pathways (53) (Fig. S4). Therefore, succinate induction is predicted to cause a decrease in metabolic flux through both the enterobactin and ubiquinone biosynthetic pathways, which should alleviate inhibition of SlyA. In view of these observations, we hypothesized that succinate induction of virulence gene expression might be occurring via SlyA. To test this hypothesis, we measured *pagC* expression in low phosphate minimal medium (LPM) with succinate as the sole carbon source and compared it to expression levels in cultures grown in the same medium with glycerol, under conditions described in the earlier study (53). Wild-type *S*. Typhimurium was compared to a *slyA* mutant strain, as well as to mutant strains complemented with either wild-type *slyA* or a mutant allele (T66A) that is insensitive to inhibition by salicylate (23). The *pagC* reporter exhibited modest succinate induction in the wild-type strain, which was abrogated by mutation of *slyA* (Fig. 4C). Complementation with *slyA* on a multi-copy vector restored induction, although expression was also increased in the glycerol control culture, likely due to gene dosage effects. Complementation with *slyA*_T66A_ abrogated the succinate-induction phenotype, indicating that induction occurs as a result of down-regulation of the enterobactin and ubiquinone pathways, which alleviates ligand-mediated allosteric inhibition of SlyA. Based on these observations, we conclude that succinate induction occurs, at least in part, through the de-repression of SlyA resulting from perturbations in aromatic carboxylate metabolism.

## DISCUSSION

The MarR transcription factor family has been characterized primarily for its role in regulating the clinically important traits of virulence and antibiotic resistance. However, its evolutionary ancestry suggests a more primordial function. All MFTFs possess small molecule binding sites allowing allosteric inhibition, but for many conserved MFTFs such as SlyA, endogenous ligands have not been identified. In this study, we show that SlyA senses perturbations in the metabolism of aromatic carboxylates, which are required for the synthesis of electron-carrying quinones and iron-scavenging catecholate siderophores, both of which are critical for bacterial respiration. Although previous studies have shown that the related MFTF, MarR, binds 2,3-DHB *in vitro* (55), no biological function was demonstrated for this interaction. We show here that SlyA can bind and respond allosterically to the ubiquinone and enterobactin precursors 4-HB and 2,3-DHB, respectively, in response to environmental and physiological cues. Metabolic flux through these pathways signals the bacterial metabolic status with respect to respiration and iron availability.

The relationship between MarR transcription factors and metabolism is further supported by the recent description of a related MFTF, CioR, in the environmental pathogen *Chromobacterium violaceum* (56). CioR regulates the iron-dependent expression of the *cioRAB* operon, which also encodes cytochrome *bd*, an iron-requiring terminal oxidase in the electron transport chain. Additional studies have identified MFTFs including YodB and CatR of *Bacillus subtilis* (57, 58), and QsrR and MhqR of *Staphylococcus aureus* (59, 60), which bind and regulate the detoxification of tail-less quinone oxidants and electrophiles to maintain cellular redox balance (61, 62).

Our observations indicate that SlyA is effectively a novel iron sensor, distinct from either Fur or its associated sRNA, RyhB (34, 63), which detects changes in the iron-dependent rate of enterobactin biosynthesis rather than sensing iron directly. Both the *entCEBAH* and *fepA-entD* operons, encoding enzymes in the enterobactin biosynthetic pathway, are co-repressed by Fur and ferrous iron and up-regulated by the catabolite repression protein CRP (64). During periods of iron limitation, apo-Fur dissociates from DNA, derepressing *ent* gene expression and inhibiting SlyA through the accumulation of enterobactin precursors. CRP can also activate the *entCEBAH* operon in response to changing carbon source availability.

The evolutionary history and broad distribution of the MFTFs suggest they have long played a central role in bacterial regulatory networks. The strong conservation of their ligand binding sites and linkage to efflux pumps (5) suggest that they evolved to sense the accumulation of potentially toxic aromatic compounds in the cell and regulate their export as a homeostatic mechanism. Small aromatic compounds can form stable complexes with metal ions and participate in redox reactions that lead to protein and DNA damage (65). These compounds can also perturb the lipid bilayer or actively transport electrons across the membrane, resulting in the disruption of membrane potential and dissipation of proton motive force {Sanchez-Maldonado, 2011; Lou, 2012}. Analogs of aromatic metabolites may act as metabolic toxins by inhibiting the biosynthesis of physiological compounds. Thus, there is an obvious fitness benefit in the ability to sense these small molecules and regulate their export from the cell (5). SlyA directly regulates the putative multidrug transporter-encoding *ydhIJK* operon, which is divergently transcribed from *slyA* in *Salmonella*. No clear function has been established for YdhIJK, and although we have previously shown that it provides resistance to exogenous fusaric acid (23), its primary function is more likely to export an unidentified endogenous compound. YdhIJK exhibits homology to the AaeAB efflux pump (34.1% and 21.1% identity between YdhJ/AaeA and YdhK/AaeB, respectively), which exports three aromatic carboxylates: 6-hydroxy-2-naphthoic acid, 2-hydroxycinnamate, and 4-HB (66). Notably, 2,3-DHB is not exported by AaeAB, but multiple pumps are able to export aromatic molecules (66). The *ydhIJK* genes do not exhibit strong genetic linkage to *slyA* throughout the *Enterobacterales* (23), suggesting that other pumps may export SlyA ligands. The physiological function of YdhIJK as well as the identity of other SlyA-associated pumps remain to be determined.

It is noteworthy that salicylate, the first well-characterized MFTF ligand (6), is a common bacterial metabolite and functional siderophore in many bacterial species (67), even though it is not synthesized by *S*. Typhimurium or *E. coli* K-12. Salicylate is also a precursor to alternative siderophores, including yersiniabactin in *Yersinia spp*. (68) and pyochelin in *Pseudomonas aeruginosa* (69). Although enterobactin is limited to Gram-negative bacterial species, catecholate siderophores are distributed throughout the bacterial kingdom (70). While few archaeal siderophores have been identified, Archaea are known to encode many homologs of catecholate siderophore synthetases that interact with 2,3-DHB, such as EntB, VibB (71), and DhbF (72), suggesting that comparable pathways may exist. Similarly, Archaea appear to synthesize 4-HB as a precursor to the electron carrier menaquinone (73, 74), and archaeal MFTFs have been shown to bind aromatic carboxylates (75). This suggests that the relationship between aromatic metabolism and MFTFs may be widespread throughout Bacteria and Archaea. Given these observations, we consider it likely that MFTFs evolved as early regulatory sensors to maintain metabolic homeostasis prior to the divergence of Bacteria and Archaea.

## MATERIALS AND METHODS

### Bacterial strains, plasmids and general reagents

All oligonucleotides, plasmids, and strains used in this study are listed in Tables S1, S2, and S3, respectively. Mutant strains were constructed in wild-type *S*. Typhimurium strain ATCC 14028s, using the λ-Red recombinase system (76). Briefly, the kanamycin cassette from pKD4 was amplified via PCR with primers encoding 40-bp flanking arms homologous to the region to be mutated and electroporated into 14028s containing pKD46 following established protocols (76). The *pagC*-*egfp* reporter plasmid pRW79 was constructed by amplifying the *pagC* promoter region using the primers PpagC-F and PpagC-R. This fragment was then ligated into pJ251-GERC following digestion with NdeI and AscI. pJ251-GERC was a gift from George Church (RRID:Addgene_47441). All cultures were grown in Luria-Bertani broth with agitation at 37°C unless otherwise indicated. Recombinant SlyA was purified as described elsewhere (77).

### RNA-Seq

Five ml LB cultures were inoculated with wildtype 14028s, *slyA*, or *tolC* mutant strains and grown overnight before a 1:500 dilution into fresh medium. Cultures were then grown to approximately 1.5 OD_600_ before centrifugation and washing three times in equal volumes of N-minimal medium (78) containing 10 µM MgSO_4_. Cultures were resuspended in N-minimal medium containing 10 µM MgSO_4_ and grown at 37°C with agitation for 1 h before 1 ml of culture was collected in 110 µl ice-cold RNA stop solution (95% v/v ethanol, 5% v/v acid-buffered phenol). Cells were pelleted by centrifugation, supernatant was discarded, and pellets stored at -80°C until the RNA was purified. RNA was purified using TRIzol (Life Technologies, Carlsbad CA) and Direct-zol miniprep columns (Zymo Research, Irvine CA) using a protocol previously described by Culviner and Laub (79). Following purification, RNA purity, integrity, and concentration were determined using a Bioanalyzer (Agilent, Santa Clara, CA) and a GE Nanovue spectrophotometer (Cytiva, Marlborough MA). Samples were depleted of rRNA using the DIY method described by Culviner and Laub (79). Depletion of rRNA was confirmed by electrophoresis of an RNA sample in a 1% agarose TAE gel containing 1% bleach (v/v) (80). RNA was then quantified on a Qubit fluorimeter (Thermo Fisher Scientific, Waltham MA). cDNA libraries were prepared from depleted RNA using NEBNext Ultra II Directional RNA Library Prep Kits and NEBNext Multiplex Oligos (New England Biolabs, Ipswich MA). Final libraries were quantified by Qubit (Thermo Fisher Scientific) and Bioanalyzer (Agilent) before submission to the Northwest Genomics Center at the University of Washington for sequencing on a NextSeq 550 (Illumina, San Diego, CA). RNA-Seq data were analyzed on the Galaxy platform (81). Reads with length or quality scores of less than 20 were discarded using Cutadapt (82). Remaining reads were aligned with the 14028s genome using Bowtie2 (83, 84). Individual gene read counts were determined using htseq-count (85), and the *slyA* and *tolC* libraries were compared to the wildtype library using DESeq2(86).

### qRT-PCR analysis of gene expression

Unless otherwise indicated, cultures were grown in LB and induced in N-minimal medium as described above. One ml of culture was added to RNA stop solution, pelleted, and RNA purified via TRIzol (Life Technologies) and Direct-zol miniprep columns (Zymo Research). RNA was quantified on a Nanovue spectrophotometer (Cytiva) and cDNA generated using the Qiagen QuantiTect Reverse Transcription kit (Qiagen, Hilden Germany). cDNA was quantified on a Bio-Rad CFX96 (Bio-Rad, Hercules CA) using SYBR Green Master Mix (87). For succinate induction experiments, cultures were grown as described by Rosenberg *et al*. (53). Five ml of LB were inoculated with each strain and incubated at room temperature for 16 h without agitation. Ampicillin was added to cultures with pWSK29-based complementation constructs to maintain the plasmid. Cultures were then diluted to 0.01 OD_600_/ml in 2 ml LPM containing either 10 mM succinate or 38 mM glycerol as the sole carbon source (53) in 13×100 mm cultures tubes. Cultures were incubated at 37°C for eight h before samples were collected for RNA purification and analysis.

### GFP reporter analysis of gene expression

Bacterial strains carrying pRW79 were grown overnight in LB and diluted into fresh medium. Cultures were grown at 37°C with shaking until they reached late exponential phase (∼1 OD_600_). Cultures were washed three times in N-minimal medium as described above and resuspended in N-minimal medium at a final concentration of 0.5 OD_600_. Cultures were aliquoted on 96-well plates along with aromatic carboxylates or 2,2-dipyridyl as indicated and incubated at 37°C for six h. Fluorescence and OD_600_ were measured every 15 min.

### NMR analysis of SlyA-ligand complexes

SlyA was overexpressed in cells grown in M9 minimal medium supplemented with ^15^N ammonium chloride and purified as previously described (23). Protein and ligand solutions were prepared in buffer containing 25 mM potassium phosphate (pH 7.0) and 150 mM NaCl. Spectra were collected from samples containing 300 µM SlyA, indicated ligand concentrations, and 8% D_2_O at 35°C on a Bruker DMX 500 Mhz spectrometer (Bruker, Billerica MA). Data were processed using NMR-Pipe (88) and NMR-View (89).

### Isothermal titration calorimetery

ITC experiments were performed at the Hans Neurath Biophysics Core (University of Washington) using an Affinity ITC (TA Instruments; New Castle DE) and analyzed using NanoAnalyze (TA Instruments). Protein and ligand solutions were prepared in buffer consisting of 50 mM potassium phosphate (pH 7.5), 100 mM NaCl, 0.1 mM EDTA, and 1 mM β-mercaptoethanol. Protein was dialyzed extensively against ITC buffer to ensure removal of contaminants, and the pH of 2,3-DHB and 4-HB solutions was confirmed before each experiment to avoid the generation of excessive background heat. Reactions were performed at 10°C with 300 µl of SlyA-containing buffer in the cell at a starting concentration of 129 µM. Ligand was then added in 2 µl injections at 4 min intervals until the reaction was saturated.

## ACKNOWLEDGMENTS

The authors would like to thank Steve Libby for helpful discussions about SlyA and its role in *S*. Typhimurium, as well Peter Brzovic and Lisa Tuttle for assistance with NMR and ITC and the Klevit lab (University of Washington) for the use of NMR facilities. ITC experiments were performed in the Hans Neurath Biophysics Core (University of Washington). The National Institutes of Health provided funding to F.C.F. (AI150041).

## REFERENCES

1. R. B. Helling et al., Toxic waste disposal in Escherichia coli. J Bacteriol 184, 3699–3703 (2002).

2. E. Pérez-Rueda, J. Collado-Vides, Common history at the origin of the position-function correlation in transcriptional regulators in archaea and bacteria. J Mol Evol 53, 172–179 (2001).

3. E. Pérez-Rueda, J. Collado-Vides, L. Segovia, Phylogenetic distribution of DNA-binding transcription factors in bacteria and archaea. Comput Biol Chem 28, 341–350 (2004).

4. A. Grove, Regulation of Metabolic Pathways by MarR Family Transcription Factors. Comput Struct Biotechnol J 15, 366–371 (2017).

5. W. R. Will, F. C. Fang, The evolution of MarR family transcription factors as counter-silencers in regulatory networks. Curr Opin Microbiol 55, 1–8 (2020).

6. S. P. Cohen, W. Yan, S. B. Levy, A multidrug resistance regulatory chromosomal locus is widespread among enteric bacteria. J Infect Dis 168, 484–488 (1993).

7. W. W. Navarre et al., Co-regulation of Salmonella enterica genes required for virulence and resistance to antimicrobial peptides by SlyA and PhoP/PhoQ. Mol Microbiol 56, 492–508 (2005).

8. W. R. Will, W. W. Navarre, F. C. Fang, Integrated circuits: how transcriptional silencing and counter-silencing facilitate bacterial evolution. Curr Opin Microbiol 23, 8–13 (2015).

9. E. A. Groisman, H. Ochman, Pathogenicity islands: bacterial evolution in quantum leaps. Cell 87, 791–794 (1996).

10. W. W. Navarre, The Impact of Gene Silencing on Horizontal Gene Transfer and Bacterial Evolution. Adv Microb Physiol 69, 157–186 (2016).

11. W. W. Navarre et al., Selective silencing of foreign DNA with low GC content by the H-NS protein in Salmonella. Science 313, 236–238 (2006).

12. I. Baglivo et al., MucR protein: Three decades of studies have led to the identification of a new H-NS-like protein. Mol Microbiol 123, 154–167 (2025).

13. 1 I. S. Barton et al., MucR acts as an H-NS-like protein to silence virulence genes and structure the nucleoid. mBio 14, e0220123 (2023).

14. B. R. Gordon, R. Imperial, L. Wang, W. W. Navarre, J. Liu, Lsr2 of Mycobacterium represents a novel class of H-NS-like proteins. J Bacteriol 190, 7052–7059 (2008).

15. B. R. Gordon et al., Lsr2 is a nucleoid-associated protein that targets AT-rich sequences and virulence genes in Mycobacterium tuberculosis. Proc Natl Acad Sci U S A 107, 5154–5159 (2010).

16. C. Tendeng, O. A. Soutourina, A. Danchin, P. N. Bertin, MvaT proteins in Pseudomonas spp.: a novel class of H-NS-like proteins. Microbiology (Reading) 149, 3047–3050 (2003).

17. P. Ding et al., A Novel AT-Rich DNA Recognition Mechanism for Bacterial Xenogeneic Silencer MvaT. PLoS Pathog 11, e1004967 (2015).

18. W. K. Smits, A. D. Grossman, The transcriptional regulator Rok binds A+T-rich DNA and is involved in repression of a mobile genetic element in Bacillus subtilis. PLoS Genet 6, e1001207 (2010).

19. D. Parke, L. N. Ornston, Hydroxycinnamate (hca) catabolic genes from Acinetobacter sp. strain ADP1 are repressed by HcaR and are induced by hydroxycinnamoyl-coenzyme A thioesters. Appl Environ Microbiol 69, 5398–5409 (2003).

20. M. Hong, M. Fuangthong, J. D. Helmann, R. G. Brennan, Structure of an OhrR-ohrA operator complex reveals the DNA binding mechanism of the MarR family. Mol Cell 20, 131–141 (2005).

21. K. J. Newberry, M. Fuangthong, W. Panmanee, S. Mongkolsuk, R. G. Brennan, Structural mechanism of organic hydroperoxide induction of the transcription regulator OhrR. Mol Cell 28, 652–664 (2007).

22. Z. Hao et al., The multiple antibiotic resistance regulator MarR is a copper sensor in Escherichia coli. Nat Chem Biol 10, 21–28 (2014).

23. W. R. Will et al., The Evolution of SlyA/RovA Transcription Factors from Repressors to Countersilencers in Enterobacteriaceae. MBio 10 (2019).

24. A. Haraga, M. B. Ohlson, S. I. Miller, Salmonellae interplay with host cells. Nat Rev Microbiol 6, 53–66 (2008).

25. J. E. Galán, R. Curtiss, Cloning and molecular characterization of genes whose products allow Salmonella typhimurium to penetrate tissue culture cells. Proc Natl Acad Sci U S A 86, 6383–6387 (1989).

26. M. Hensel et al., Simultaneous identification of bacterial virulence genes by negative selection. Science 269, 400–403 (1995).

27. H. Ochman, F. C. Soncini, F. Solomon, E. A. Groisman, Identification of a pathogenicity island required for Salmonella survival in host cells. Proc Natl Acad Sci U S A 93, 7800–7804 (1996).

28. J. E. Shea, M. Hensel, C. Gleeson, D. W. Holden, Identification of a virulence locus encoding a second type III secretion system in Salmonella typhimurium. Proc Natl Acad Sci U S A 93, 2593–2597 (1996).

29. C. M. Alpuche Aranda, J. A. Swanson, W. P. Loomis, S. I. Miller, Salmonella typhimurium activates virulence gene transcription within acidified macrophage phagosomes. Proc Natl Acad Sci U S A 89, 10079–10083 (1992).

30. A. K. Heroven, G. Nagel, H. J. Tran, S. Parr, P. Dersch, RovA is autoregulated and antagonizes H-NS-mediated silencing of invasin and rovA expression in Yersinia pseudotuberculosis. Mol Microbiol 53, 871–888 (2004).

31. M. M. Haque, M. S. Kabir, L. Q. Aini, H. Hirata, S. Tsuyumu, SlyA, a MarR family transcriptional regulator, is essential for virulence in Dickeya dadantii 3937. J Bacteriol 191, 5409–5418 (2009).

32. N. Weatherspoon-Griffin, H. J. Wing, Characterization of SlyA in Shigella flexneri Identifies a Novel Role in Virulence. Infect Immun 84, 1073–1082 (2016).

33. M. Palacios et al., Identification of Two Regulators of Virulence That Are Conserved in Klebsiella pneumoniae Classical and Hypervirulent Strains. MBio 9 (2018).

34. G. Porcheron, C. M. Dozois, Interplay between iron homeostasis and virulence: Fur and RyhB as major regulators of bacterial pathogenicity. Vet Microbiol 179, 2–14 (2015).

35. H. Nikaido, Multidrug resistance in bacteria. Annu Rev Biochem 78, 119–146 (2009).

36. E. A. Groisman, A. Duprey, J. Choi, How the PhoP/PhoQ System Controls Virulence and Mg2+ Homeostasis: Lessons in Signal Transduction, Pathogenesis, Physiology, and Evolution. Microbiol Mol Biol Rev 85, e0017620 (2021).

37. R. Bentley, The shikimate pathway--a metabolic tree with many branches. Crit Rev Biochem Mol Biol 25, 307–384 (1990).

38. V. V. Shende, K. D. Bauman, B. S. Moore, The shikimate pathway: gateway to metabolic diversity. Nat Prod Rep 41, 604–648 (2024).

39. R. Meganathan, Ubiquinone biosynthesis in microorganisms. FEMS Microbiol Lett 203, 131–139 (2001).

40. K. N. Raymond, E. A. Dertz, S. S. Kim, Enterobactin: an archetype for microbial iron transport. Proc Natl Acad Sci U S A 100, 3584–3588 (2003).

41. S. C. Andrews, A. K. Robinson, F. Rodríguez-Quiñones, Bacterial iron homeostasis. FEMS Microbiol Rev 27, 215–237 (2003).

42. E. R. Frawley, F. C. Fang, The ins and outs of bacterial iron metabolism. Mol Microbiol 93, 609–616 (2014).

43. J. Lawrence, G. B. Cox, F. Gibson, Biosynthesis of ubiquinone in Escherichia coli K-12: biochemical and genetic characterization of a mutant unable to convert chorismate into 4-hydroxybenzoate. J Bacteriol 118, 41–45 (1974).

44. I. G. Young, R. A. Leppik, J. A. Hamilton, F. Gibson, Biochemical and genetic studies on ubiquinone biosynthesis in Escherichia coli K-12:4-hydroxybenzoate octaprenyltransferase. J Bacteriol 110, 18–25 (1972).

45. R. K. Luke, F. Gibson, Location of three genes concerned with the conversion of 2,3-dihydroxybenzoate into enterochelin in Escherichia coli K-12. J Bacteriol 107, 557–562 (1971).

46. I. G. Young, L. Langman, R. K. Luke, F. Gibson, Biosynthesis of the iron-transport compound enterochelin: mutants of Escherichia coli unable to synthesize 2,3-dihydroxybenzoate. J Bacteriol 106, 51–57 (1971).

47. C. Dahm, R. Müller, G. Schulte, K. Schmidt, E. Leistner, The role of isochorismate hydroxymutase genes entC and menF in enterobactin and menaquinone biosynthesis in Escherichia coli. Biochim Biophys Acta 1425, 377–386 (1998).

48. D. J. Shaw, E. C. Robinson, R. Meganathan, R. Bentley, J. R. Guest, Recombinant plasmids containing menaquinone biosynthetic genes of Escherichia coli. FEMS Microbiology Letters 17, 63–67 (1983).

49. S. J. Taylor, S. E. Winter, Salmonella finds a way: Metabolic versatility of Salmonella enterica serovar Typhimurium in diverse host environments. PLoS Pathog 16, e1008540 (2020).

50. B. D. Bennett et al., Absolute metabolite concentrations and implied enzyme active site occupancy in Escherichia coli. Nat Chem Biol 5, 593–599 (2009).

51. A. Bagg, J. B. Neilands, Ferric uptake regulation protein acts as a repressor, employing iron (II) as a cofactor to bind the operator of an iron transport operon in Escherichia coli. Biochemistry 26, 5471–5477 (1987).

52. L. Escolar, J. Pérez-Martín, V. de Lorenzo, Opening the iron box: transcriptional metalloregulation by the Fur protein. J Bacteriol 181, 6223–6229 (1999).

53. G. Rosenberg et al., Host succinate is an activation signal for Salmonella virulence during intracellular infection. Science 371, 400–405 (2021).

54. 1 L. Spiga et al., An Oxidative Central Metabolism Enables Salmonella to Utilize Microbiota-Derived Succinate. Cell Host Microbe 22, 291-301.e296 (2017).

55. L. M. Chubiz, C. V. Rao, Aromatic acid metabolites of Escherichia coli K-12 can induce the marRAB operon. J Bacteriol 192, 4786–4789 (2010).

56. B. B. Batista, V. M. de Lima, W. R. Will, F. C. Fang, J. F. da Silva Neto, A cytochrome bd repressed by a MarR family regulator confers resistance to metals, nitric oxide, sulfide, and cyanide in Chromobacterium violaceum. Appl Environ Microbiol 91, e0236024 (2025).

57. B. K. Chi et al., The redox-sensing regulator YodB senses quinones and diamide via a thiol-disulfide switch in Bacillus subtilis. Proteomics 10, 3155–3164 (2010).

58. S. J. Lee et al., Two distinct mechanisms of transcriptional regulation by the redox sensor YodB. Proc Natl Acad Sci U S A 113, E5202–5211 (2016).

59. Q. Ji et al., Molecular mechanism of quinone signaling mediated through S-quinonization of a YodB family repressor QsrR. Proc Natl Acad Sci U S A 110, 5010–5015 (2013).

60. V. N. Fritsch et al., The MarR-Type Repressor MhqR Confers Quinone and Antimicrobial Resistance in Staphylococcus aureus. Antioxid Redox Signal 31, 1235–1252 (2019).

61. T. Franza, P. Gaudu, Quinones: more than electron shuttles. Res Microbiol 173, 103953 (2022).

62. M. Liebeke et al., Depletion of thiol-containing proteins in response to quinones in Bacillus subtilis. Mol Microbiol 69, 1513–1529 (2008).

63. E. Massé, S. Gottesman, A small RNA regulates the expression of genes involved in iron metabolism in Escherichia coli. Proc Natl Acad Sci U S A 99, 4620–4625 (2002).

64. X. Gu, Z. Zhang, W. Huang, Rapid evolution of expression and regulatory divergences after yeast gene duplication. Proc Natl Acad Sci U S A 102, 707–712 (2005).

65. N. Schweigert, A. J. Zehnder, R. I. Eggen, Chemical properties of catechols and their molecular modes of toxic action in cells, from microorganisms to mammals. Environ Microbiol 3, 81–91 (2001).

66. T. K. Van Dyk, L. J. Templeton, K. A. Cantera, P. L. Sharpe, F. S. Sariaslani, Characterization of the Escherichia coli AaeAB efflux pump: a metabolic relief valve? J Bacteriol 186, 7196–7204 (2004).

67. C. Ratledge, L. P. Macham, K. A. Brown, B. J. Marshall, Iron transport in Mycobacterium smegmatis: a restricted role for salicylic acid in the extracellular environment. Biochim Biophys Acta 372, 39–51 (1974).

68. R. D. Perry, P. B. Balbo, H. A. Jones, J. D. Fetherston, E. DeMoll, xYersiniabactin from Yersinia pestis: biochemical characterization of the siderophore and its role in iron transport and regulation. Microbiology (Reading) 145 (Pt 5), 1181–1190 (1999).

69. C. D. Cox, K. L. Rinehart, M. L. Moore, J. C. Cook, Pyochelin: novel structure of an iron-chelating growth promoter for Pseudomonas aeruginosa. Proc Natl Acad Sci U S A 78, 4256–4260 (1981).

70. J. Kramer, Ö. Özkaya, R. Kümmerli, Bacterial siderophores in community and host interactions. Nat Rev Microbiol 18, 152–163 (2020).

71. T. A. Keating, C. G. Marshall, C. T. Walsh, Reconstitution and characterization of the Vibrio cholerae vibriobactin synthetase from VibB, VibE, VibF, and VibH. Biochemistry 39, 15522–15530 (2000).

72. J. J. May, T. M. Wendrich, M. A. Marahiel, The dhb operon of Bacillus subtilis encodes the biosynthetic template for the catecholic siderophore 2,3-dihydroxybenzoate-glycine-threonine trimeric ester bacillibactin. J Biol Chem 276, 7209–7217 (2001).

73. B. Nowicka, J. Kruk, Occurrence, biosynthesis and function of isoprenoid quinones. Biochim Biophys Acta 1797, 1587–1605 (2010).

74. M. Lübben, Cytochromes of archaeal electron transfer chains. Biochim Biophys Acta 1229, 1–22 (1995).

75. V. Duval, L. M. McMurry, K. Foster, J. F. Head, S. B. Levy, Mutational analysis of the multiple-antibiotic resistance regulator MarR reveals a ligand binding pocket at the interface between the dimerization and DNA binding domains. J Bacteriol 195, 3341–3351 (2013).

76. K. A. Datsenko, B. L. Wanner, One-step inactivation of chromosomal genes in Escherichia coli K-12 using PCR products. Proc Natl Acad Sci U S A 97, 6640–6645 (2000).

77. W. R. Will, D. H. Bale, P. J. Reid, S. J. Libby, F. C. Fang, Evolutionary expansion of a regulatory network by counter-silencing. Nat Commun 5, 5270 (2014).

78. M. D. Snavely, C. G. Miller, M. E. Maguire, The mgtB Mg2+ transport locus of Salmonella typhimurium encodes a P-type ATPase. J Biol Chem 266, 815–823 (1991).

79. P. H. Culviner, C. K. Guegler, M. T. Laub, A Simple, Cost-Effective, and Robust Method for rRNA Depletion in RNA-Sequencing Studies. mBio 11 (2020).

80. P. S. Aranda, D. M. LaJoie, C. L. Jorcyk, Bleach gel: a simple agarose gel for analyzing RNA quality. Electrophoresis 33, 366–369 (2012).

81. G. Community, The Galaxy platform for accessible, reproducible, and collaborative data analyses: 2024 update. Nucleic Acids Res 52, W83–W94 (2024).

82. M. Marcel, Cutadapt removes adapter sequences from high-throughput sequencing reads. EMBnet.journal 17, 10–12 (2011).

83. B. Langmead, C. Trapnell, M. Pop, S. L. Salzberg, Ultrafast and memory-efficient alignment of short DNA sequences to the human genome. Genome Biol 10, R25 (2009).

84. B. Langmead, S. L. Salzberg, Fast gapped-read alignment with Bowtie 2. Nat Methods 9, 357–359 (2012).

85. S. Anders, P. T. Pyl, W. Huber, HTSeq--a Python framework to work with high-throughput sequencing data. Bioinformatics 31, 166–169 (2015).

86. M. I. Love, W. Huber, S. Anders, Moderated estimation of fold change and dispersion for RNA-seq data with DESeq2. Genome Biol 15, 550 (2014).

87. O. Aparicio et al., Chromatin immunoprecipitation for determining the association of proteins with specific genomic sequences in vivo. Curr Protoc Mol Biol Chapter 21, Unit 21.23 (2005).

88. F. Delaglio et al., NMRPipe: a multidimensional spectral processing system based on UNIX pipes. J Biomol NMR 6, 277–293 (1995).

89. B. A. Johnson, R. A. Blevins, NMR View: A computer program for the visualization and analysis of NMR data. J Biomol NMR 4, 603–614 (1994).

