## Supplementary Material for "The MarR Family Transcription Factor SlyA Senses Iron and Respiratory Status in Enteric Bacteria"


Contents:


Supplementary Figure S1. Exogenous aromatic metabolites inhibit *pagC* expression.

Supplementary Figure S2. Dose-dependent conformational changes in SlyA upon ligand binding.

Supplementary Figure S3. Salicylate binds SlyA and induces conformational changes.

Supplementary Figure S4. Succinate increases the expression of aromatic carboxylate metabolism genes.

Supplementary Figure S5. A general model for the regulation of SlyA by aromatic carboxylate metabolism.

Supplementary Table S1. Oligonucleotides used in this study.

Supplementary Table S2. Plasmids used in this study.

Supplementary Table S3. Strains used in this study.

Supplementary References.


Analyzed RNA-Seq data are attached separately:

Supplementary Dataset S1. Misregulated genes in *slyA.*

Supplementary Dataset S2. Misregulated genes in *tolC.*

RNA-Seq datasets are available on the Gene Expression Omnibus (GSE 293030; https://www.ncbi.nlm.nih.gov/geo/query/acc.cgi?acc=GSE293030);

### Supplementary Figures.

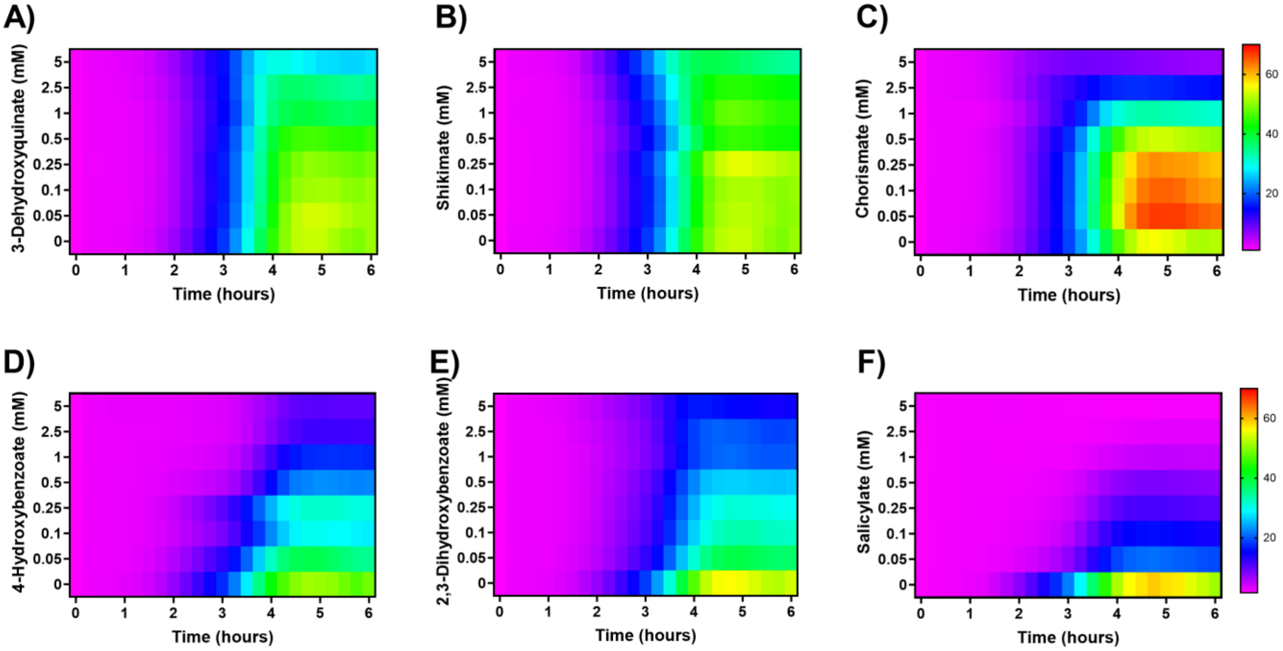


**Figure S1. Exogenous aromatic metabolites inhibit *pagC* expression.** To determine whether specific aromatic metabolites inhibit *pagC* expression, *S.* Typhimurium carrying a *pagC-egfp* transcriptional fusion was grown at 37ºC in N-minimal medium containing 10 µM $MgSO_4$ in the presence of increasing concentrations of 3-dehydroquinate (A), shikimate (B), chorismate (C), 4-hydroxybenzoate (D), 2,3-dehydroxybenzoate (E), and salicylate (F) (indicated on the y-axis). Quantities are indicated by scale bars on the right. Fluorescence was measured at 15 min intervals and normalized to the t=0 sample. Data represent the mean of three independent experiments.

A)

B)

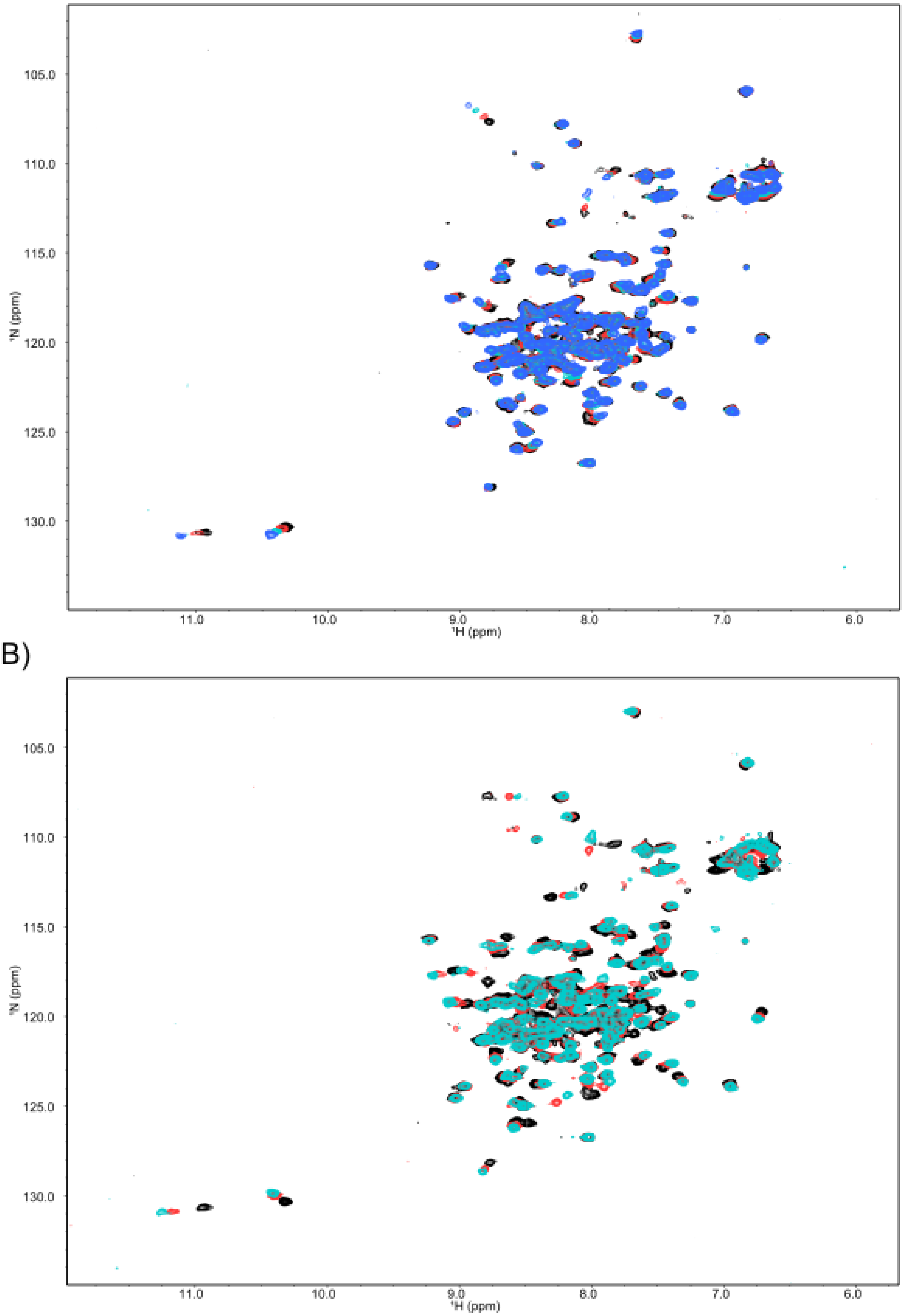


**Figure S2. Dose-dependent conformational changes in SlyA upon ligand binding.** To determine whether the spectra shown in Fig. 3., $^{1}H$,$^{15}N$-HSQC NMR spectroscopy was performed in the presence of increasing ligand:SlyA ratios. 300 μM uniformly labelled $^{15}N$-SlyA was incubated with 4-HB (A) at 2:1 (red), 4:1 (green), and 8:1 (blue) ratios, and 2,3-DHB (B) at 2:1 (red) and 4:1 (cyan) ratios. Ligand-bound spectra were overlaid on the ligand-free apo-SlyA spectrum (black).

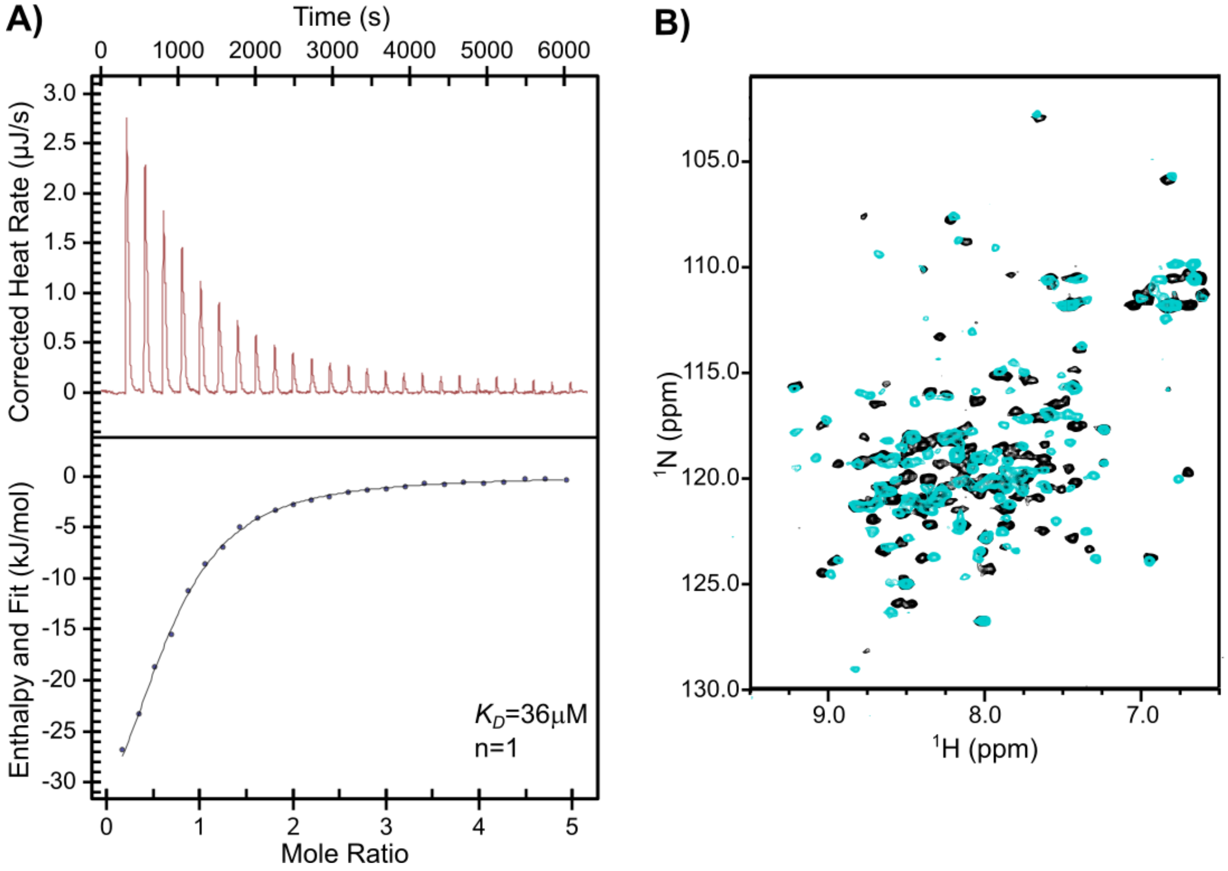


**Figure S3. Salicylate binds SlyA and induces conformational changes.** (A) ITC was performed to determine the affinity of SlyA binding salicylate. Binding reactions were performed in triplicate at 10ºC with aliquots of ligand injected at 4 min intervals. The upper panels are thermographs, measuring the heat generated following each ligand injection. The enthalpy and stoichiometry of each injection are shown in the bottom plot. Each experiment was performed three times, with representative data shown. (B) Structural changes induced by binding were observed using $^1$H,$^{15}$N-HSQC NMR spectroscopy performed on uniformly labelled $^{15}$N-SlyA in the presence (cyan) or absence (black) of salicylate. Salicylate was added to SlyA at a 4:1 molar ratio (1.2 mM:300 μM) and incubated at 35ºC.

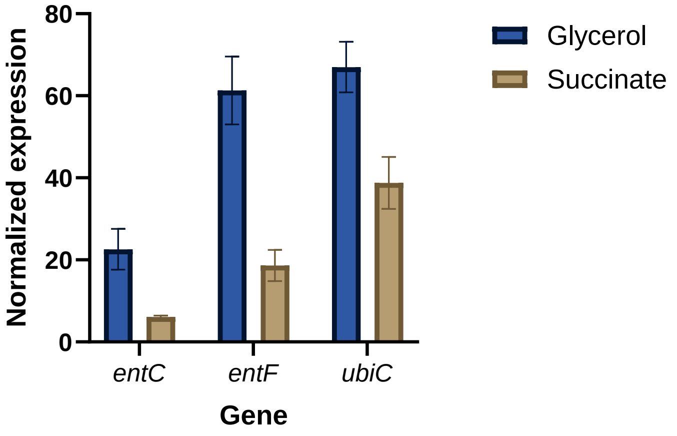


**Figure S4. Succinate increases the expression of aromatic carboxylate metabolism genes.** A published study examined the effect of succinate induction on *S.* Typhimurium virulence by performing RNA-Seq on cells grown in LPM containing either glycerol or succinate as the sole carbon source (1). Aromatic metabolism genes exhibiting a significant decrease in expression (P-value ≤ 0.005) in that study are depicted here with the author's permission.

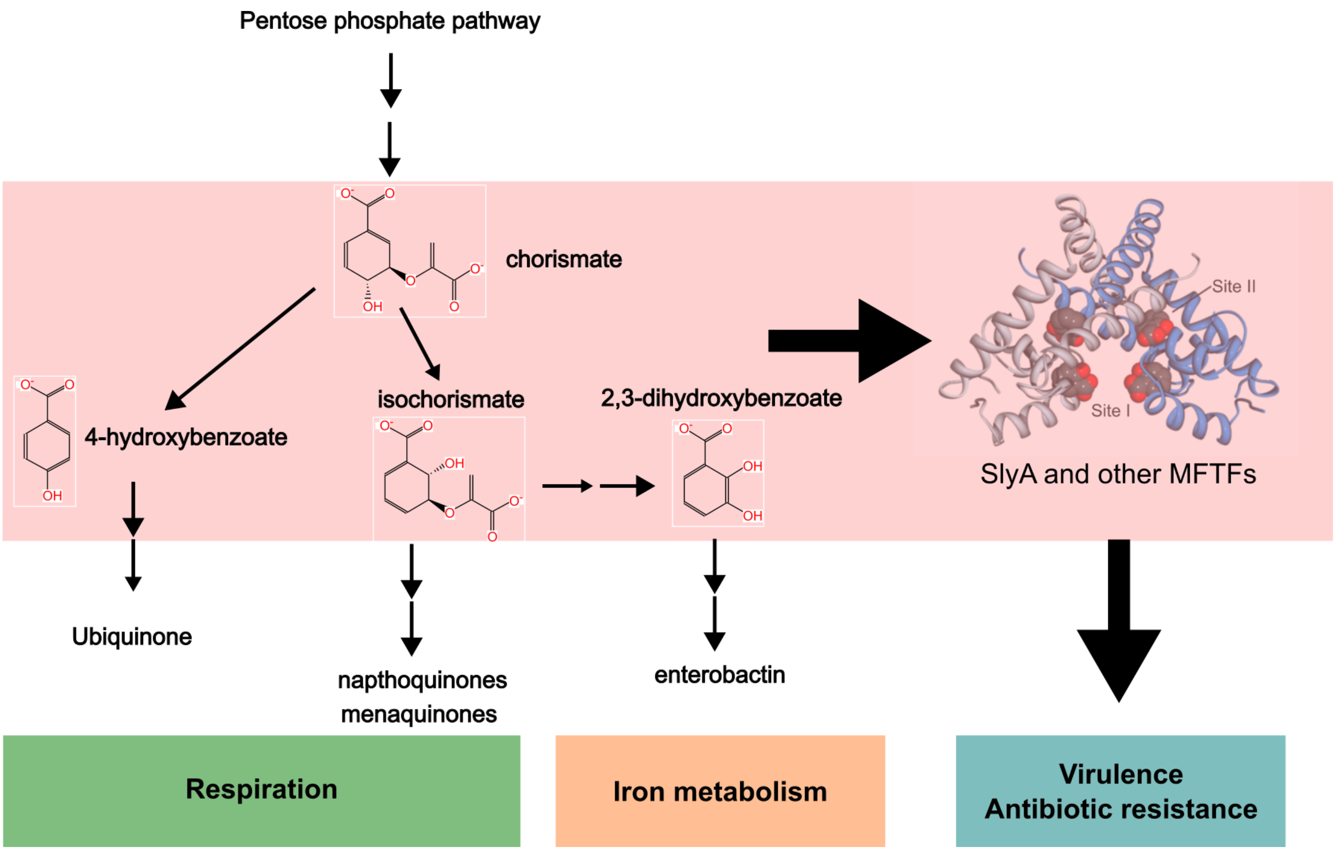


**Figure S5. A general model for the regulation of SlyA by aromatic carboxylate metabolism**. The aromatic carboxylate intermediate chorismate is synthesized via the pentose phosphate pathway. Chorismate is converted to 4-hydroxybenzoate and isochorismate as intermediates in the biosynthesis of the electron carriers ubiquinone and napthoquinone, which are required for bacterial respiration. Chorismate is also converted to 2,3-dihydroxybenzoate, a precursor to the catecholate siderophore enterobactin, which scavenges extracellular iron. Changes in either iron availability or respiration also change metabolic flux through the quinone and enterobactin biosynthetic pathways and concomitantly modulate the activity of SlyA and possibly other MFTFs to regulate virulence and antibiotic resistance.

Table S1. Oligonucleotides used in this study.

| Name | Sequence (5'-3') |
|---|---|
| aroC-kan-F | ATTAAAACACGCAAACGACAACAACGATAACGGAGCCGTGGTGTAG GCTGGAGCTGC |
| aroC-kan-R | ACCATGCCAGCAGCGCAATCGCGGTTTTTTTCATTTCTTAACATAT GAATATCCTCCTT |
| aroD-kan-F | TAAAATTATAATTGCGACGAATGACAATGAAGGGTACCAAGTGTAGG CTGGAGCTGC |
| aroD-kan-R | GTGGCAGAAAGAGAATATTCCGCCACACGATAAAGTATTACATATGA ATAT CCTCCTT |
| entC-kan-F | AAGATGAAGT GTATATAAGC CTTTATCATT GGAGGATGATGT GTA GGC TGG AGC TGC |
| entC-kan-R | CCGGCCAACGGGTGAAAGGTATACGCATCATCGTTCCTTAACATATG AATATCCTCCT T |
| entD-kan-F | ACCTTCCCTCCCTCATTCGGGGAGGGAATTGGCAAAAACGGTGTAG GCTGGAGCTGC |
| entD-kan-R | AGAAACGTGAAAATCATTCAGCGCCATAGGGATCTCATTTAACATATG AA TATCCTCCTT |
| menD-kan-F | TCTTTATACTTAGTCCCAATGATTGATACCGGACAAACTCGTGTAGG CTGGAGCTGC |
| menD-kan-R | GTCCCAGGCTGTCCGGGCATGTGCTGCGCGTGCAACATCAACATAT GAATATCCTCCTT |
| menF-kan-F | TACCCCGTATAATGTGGGGTTTTTAACAGGGAGGGTCCGCGT GTAGGCTGGAGCTGC |
| menF-kan-R | AATGGATTATTGATATGGGTCGGGAATATGTGACTCATTAACATATGA A TATCCTCCTT |
| pabA-kan-F | ACTGAGTAAAATAGTGCGGTTCTACTCACCCGGAGCCGCCGTGTAG GCTGGAGCTGCTTC |
| pabA-kan-R | AAATATGAATAAAAAATCACTCAATAGCAACCACAAATCATCCATATGA ATATCCTCCTT |
| pagC-3'-F | AGAACATTCCACTCAGGATGGCGA |
| pagC-3'-R | GACGACGATATTCTCCAGCGGATT |
| pheA-kan-F | ATCGGGGGGCCTTTTTTATTGATAACAAAAAGGCAACACTGTGTAGG CTGGAGCTGC |
| pheA-kan-R | CAGTGCCGGATGATTCACATCATCCGGCACCTTTTCATCAACATATG AATATCCTCCTT |
| PpagC-F | AGGCGCGCCGTAATGACCA AAGCATAAAAGCATG |
| PpagC-R | AACAACTCCTTAATACTACTTATTATTTACG |
| rpoD-F | GTGAATGGGCACTGTTGAACTG |
| rpoD-R | TTCCAGCAGATAGGTAATGGCTTC |
| slyA-kan-F | AGC ATAATAACTT AGCAAGCTAA TTATAAGGAG ATGAAAT GTGTAGGCTGGAGCTGC |
| slyA-kan-R | ACGTGTGGTCACATGGCCACACGTATGCCCCTGCACCTCAAACATA TGAATATCCTCCTT |

| | |
|---|---|
| tolC-kan-F | TACAAATTGA TCAGCGCTAA ATACTGCTTC ACAACAAGGA GTGTAGGCTGGAGCTGCTTC |
| tolC-kan-R | CACAGGTCTGATAAGCGCAGCGCCAGCGAATAACTTATCACATATGA ATATCCTCCTTAG |
| trpD-kan-F | GTGCTATCGC CACCGCGCAT CATGCACAGG AGACCTTCTGGT GTA GGC TGGAGCTGC |
| trpD-kan-R | TGTCTGCGACGATTTTCGCTAAAACGGTTTGCATTATTTAACATATGA ATATCCTCCTT |
| tyrA-kan-F | GAGCGGCCAGCTGGCGGTGCGCGTCGCATAAGAGGTTGTTGTGTA GGCTGGAGCTGC |
| tyrA-kan-R | AAGCCAGCAAAGCTGGCTTTTAGTATAGATGTCATCATTAACATATGA ATATCCTCCTT |
| ubiA-kan-F | TTTTTACCTGCATCGCCGCTGTACTGAGAGGAAGATAAAGGTGTAG GCTGGAGCTGC |
| ubiA-kan-R | ATTTTGGCTTTGTAGGCCGGGTCCGCCCGGCATGACATCAACATAT GAATATCCTCCTT |
| ubiC-kan-F | GAGATACAATGACTTTAGGTTATGAATCGGAGAGTAAGGCGTGTAGG CTGGAGCTGC |
| ubiC-kan-R | ACTCTGCGTCAGACTCCACTCCATCTTTATCTTCCTCTCAACATATGA ATATCCTCCTT |
| ubiE-kan-F | TACACTTCTTGAACATTTTTATCGATAAGCAGGCACTGAGGTGTAGG CTGGAGCTGC |
| ubiE-kan-R | CTGCGGTCACTAAGGGTTTAAAAGGCATTCCACCCTCCTAACATATG AATATCCTCCTT |

Table S2. Plasmids used in this study.

| **Name** | **Description** | **Source** |
|---|---|---|
| pJ251-GERC | eGFP fluorescent reporter vector | A gift from George Church; Addgene.org plasmid #47441 |
| pKD4 | Recombineering plasmid encoding kanamycin resistance cassette | (2) |
| pKD46 | λ-Red recombinase expression plasmid | (2) |
| pRW79 | pJ251-GERC *pagC-egfp* | This study |
| pSL2143 | pWSK29 *slyA* | (3) |
| pSL2143-T66A | pWSK29 *slyA* T66A | (3) |
| pWSK29 | Low copy number vector | (4) |

Table S3. Strains used in this study.

| Name | Description | Source or reference |
|---|---|---|
| 14028s | Wildtype *S. enterica* serovar Typhimurium | Fang lab collection; ATCC |
| *aroC* | 14028s Δ*aroC::kan* constructed using aroC-kan-F and aroC-kan-R | This study |
| *aroD* | 14028s Δ*aroD::kan* constructed using aroD-kan-F and aroD-kan-R | This study |
| *entC* | 14028s Δ*entC::kan* constructed using entC-kan-F and entC-kan-R | This study |
| *entD* | 14028s Δ*entD::kan* constructed using entD-kan-F and entD-kan-R | This study |
| *menD* | 14028s Δ*menD::kan* constructed using menD-kan-F and menD-kan-R | This study |
| *menF* | 14028s Δ*menF::kan* constructed using menF-kan-F and menF-kan-R | This study |
| *pabA* | 14028s Δ*pabA::kan* constructed using pabA-kan-F and pabA-kan-R | This study |
| *pheA* | 14028s Δ*pheA::kan* constructed using pheA-kan-F and pheA-kan-R | This study |
| *slyA* | 14028s Δ*slyA::kan* contstructed using slyA-kan-F and slyA-kan-R | This study |
| *tolC* | 14028s Δ*tolC::kan* constructed using tolC-kan-F and tolC-kan-R | This study |
| *trpD* | 14028s Δ*trpD::kan* constructed using trpD-kan-F and trpD-kan-R | This study |
| *tyrA* | 14028s Δ*tyrA::kan* constructed using tyrA-kan-F and tyrA-kan-R | This study |
| *ubiA* | 14028s Δ*ubiA::kan* constructed using ubiA-kan-F and ubiA-kan-R | This study |
| *ubiC* | 14028s Δ*ubiC::kan* constructed using ubiC-kan-F and ubiC-kan-R | This study |
| *ubiE* | 14028s Δ*ubiE::kan* constructed using ubiE-kan-F and ubiE-kan-R | This study |

**Supplementary References**

1. G. Rosenberg *et al.*, Host succinate is an activation signal for *Salmonella* virulence during intracellular infection. *Science* **371**, 400-405 (2021).
2. K. A. Datsenko, B. L. Wanner, One-step inactivation of chromosomal genes in *Escherichia coli* K-12 using PCR products. *Proc Natl Acad Sci U S A* **97**, 6640-6645 (2000).
3. W. R. Will *et al.*, The Evolution of SlyA/RovA Transcription Factors from Repressors to Countersilencers in *Enterobacteriaceae*. *MBio* **10** (2019).
4. R. F. Wang, S. R. Kushner, Construction of versatile low-copy-number vectors for cloning, sequencing and gene expression in *Escherichia coli*. *Gene* **100**, 195-199 (1991).
